# An antisense RNA regulates production of DnaA and affects sporulation in *Bacillus subtilis*

**DOI:** 10.1101/2025.02.18.638791

**Authors:** Emma L. Sedivy, Janet L. Smith, Alan D. Grossman

## Abstract

DnaA is the replication initiator and a transcription factor in virtually all bacteria. Although the synthesis and activity of DnaA are highly regulated, the mechanisms of regulation vary between organisms. We found that production of DnaA in *Bacillus subtilis* is regulated by an antisense RNA that overlaps with the 5’ untranslated region upstream of the *dnaA* open reading frame. We initially observed this RNA in in vitro transcription experiments and found that its production was inhibited by DnaA. This RNA, now called ArrA for **<u>a</u>**ntisense **<u>R</u>**NA **<u>r</u>**epressor of *dna**<u>A</u>***, is made in vivo. We identified the *arrA* promoter and made a mutation that greatly reduced (or eliminated) production of ArrA RNA in vitro and in vivo. In vivo, this *arrA* promoter mutation caused an increase in the amount of mRNA and protein from *dnaA* and *dnaN*, indicating that *arrA* expression normally inhibits expression of the *dnaA-dnaN* operon. The *arrA* mutation also caused a delay in sporulation that was alleviated by loss of *sda*, a sporulation-inhibitory gene that is directly activated by DnaA. *arrA* appears to be conserved in some members of the *Bacillus* genus, indicating that *arrA* has evolved in at least some endospore-forming bacteria to modulate production of DnaA and enable timely and robust sporulation.

**Author summary:** DnaA is the highly conserved replication initiator and transcription factor found in virtually all bacteria. The synthesis and activity of DnaA are highly regulated, and different types of bacteria use different mechanisms to control this key protein. We found that DnaA production in *Bacillus subtilis* is inhibited by an antisense RNA that overlaps with the 5’ untranslated region of the *dnaA* mRNA. In the absence of this antisense RNA, there was an increase in the amount of *dnaA* mRNA and protein. There was also a delay in sporulation that depends on *sda*, a sporulation-inhibitory gene that is directly activated by DnaA. *arrA* appears to be conserved in several members of the *Bacillus* genus, indicating that it has evolved to temper production DnaA and enable robust sporulation in these species.

## Introduction

DnaA is required for the initiation of DNA replication in virtually all bacteria, and also functions as a transcription factor {reviewed in: (Kaguni, 2006; Menikpurage *et al*., 2021)}. DnaA binds to sequences in the origin of replication (*oriC*) and promotes DNA unwinding and subsequent assembly of the replisome {reviewed in: (Mott and Berger, 2007; Chodavarapu and Kaguni, 2016; Rashid and Berger, 2025)}. The activity of DnaA in replication initiation and at *oriC* is highly regulated, although many of the mechanisms of regulation vary between different bacterial species (Jameson and Wilkinson, 2017; Felletti *et al*., 2019).

In most species, including *B. subtilis* and *E. coli*, *dnaA* is immediately upstream of and co-transcribed with *dnaN* (encoding the sliding clamp, a.k.a., the β-clamp, of DNA polymerase) (Fig 1A). DnaA represses transcription of the *dnaA-dnaN* operon, creating a homeostatic feedback loop that is one of the more highly conserved mechanisms in bacteria to regulate expression of *dnaA* {(Menikpurage *et al*., 2021) and references therein**}**.

**Figure 1.**
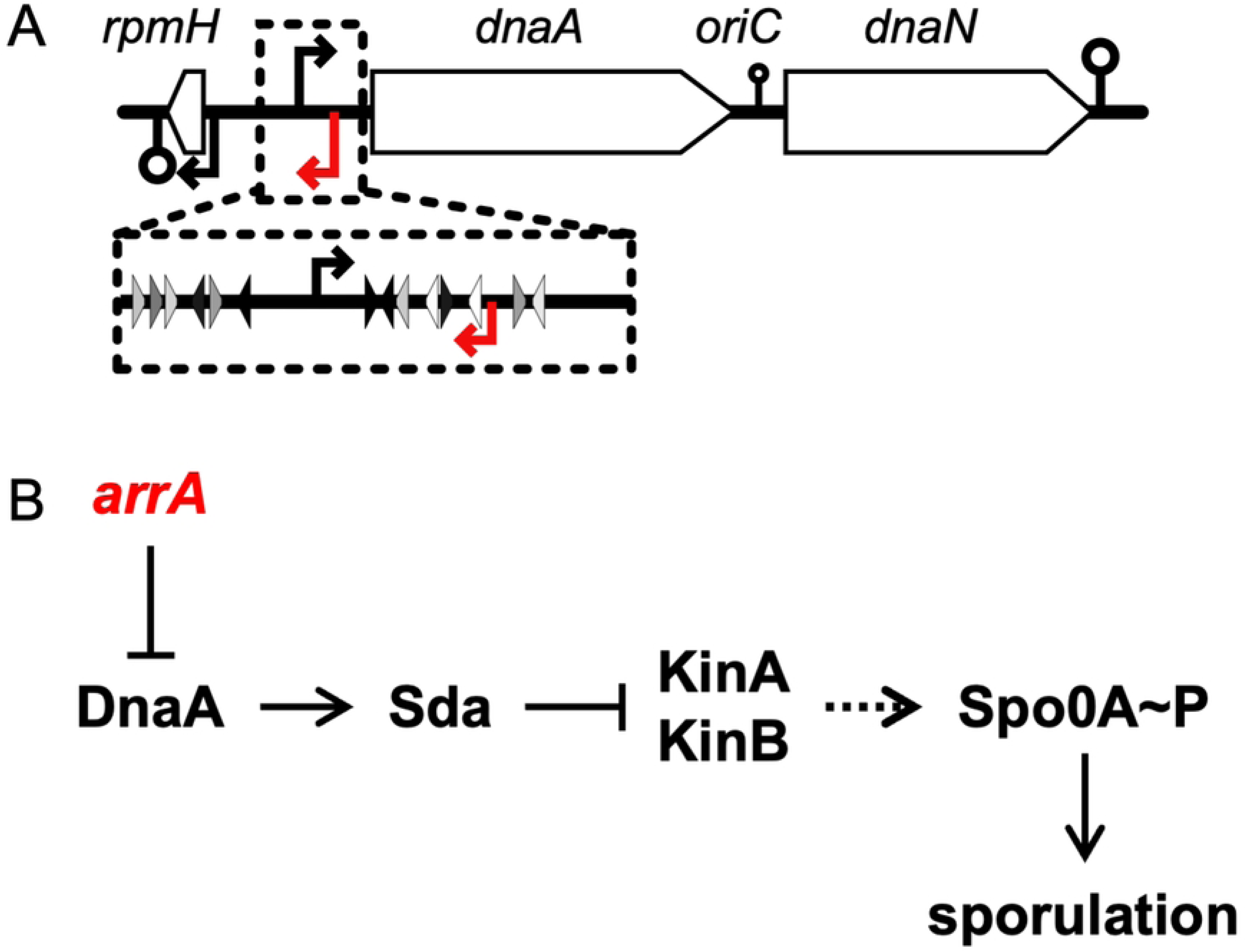
Map of the *oriC* region of the chromosome and effects of *arrA* on DnaA and sporulation. **A. Map of the *oriC* region of the *B. subtilis* chromosome.** Distances are shown approximately to scale. The black horizontal line represents the chromosome and *rpmH*, *dnaA*, and *dnaN* are indicated as white rectangles with arrows indicating the direction of transcription. Transcription start sites and terminators are indicated by arrows and stem-loops (⚲), respectively, above and below the chromosome. The red arrows indicate the transcription start site of *arrA*. The transcription terminator between *dnaA* and *dnaN* allows partial readthrough, as represented by a smaller stem-loop (⚲) symbol. Inset: The DnaA binding sites upstream of *dnaA* are indicated with small triangles showing location and direction. Black triangles indicate consensus DnaA binding sites; lighter shaded triangles are proportionally less similar to the consensus DnaA binding motif (Smith and Grossman, 2015). **B. Pathway connecting *arrA* to Spo0A and the initiation of sporulation.** The transcription factor Spo0A serves as the master regulator for entry into sporulation. Spo0A is activated by phosphorylation via a phosphorelay that involves transfer of phosphate from histidine protein kinases (primarily KinA and KinB, although others can also contribute) to Spo0F, then from Spo0F to Spo0B (not indicated), and finally to Spo0A. Spo0A∼P is essential for the initiation of sporulation and activates several genes including the sporulation genes *spoIIA, spoIIE*, and *spoIIG*. DnaA directly activates transcription of *sda,* whose product inhibits the kinases (KinA and KinB) needed for the activation of Spo0A and the initiation of sporulation. We found that transcription of *arrA* serves to decrease expression of *dnaA*, and that this function of *arrA* is crucial for the proper timing of sporulation.

DnaA also binds sequences in other promoter regions and in many cases directly modulates transcription of the associated genes {reviewed in (Menikpurage *et al*., 2021)}. In *B. subtilis*, eight promoter regions are strongly bound by DnaA (Goranov *et al*., 2005; Ishikawa *et al*., 2007; Cho *et al*., 2008; Breier and Grossman, 2009; Hoover *et al*., 2010; Smith and Grossman, 2015; Seid *et al*., 2016) and there are many transcription units whose expression is affected both directly and indirectly. By virtue of its effects on gene expression, DnaA regulates sporulation, competence development, and the response to DNA damage (Burkholder *et al*., 2001; Goranov *et al*., 2005; Hoover *et al*., 2010; Washington *et al*., 2017).

DnaA can inhibit the initiation of sporulation by activating transcription of the sporulation inhibitory gene *sda* (Burkholder *et al*., 2001; Ishikawa *et al*., 2007; Breier and Grossman, 2009). The *sda* gene product inhibits protein kinases needed for the initiation of sporulation (Burkholder *et al*., 2001; Rowland *et al*., 2004; Whitten *et al*., 2007; Cunningham and Burkholder, 2009) (Fig 1B). In this way, when there is an increase in the amount of active DnaA (e.g., when there is replication arrest), there is a subsequent increase in transcription of *sda* and sporulation initiation is inhibited (Burkholder *et al*., 2001; Rowland *et al*., 2004; Veening *et al*., 2009; Hoover *et al*., 2010).

Here we report that a small RNA (sRNA) encoded antisense to the 5’ end of the *dnaA* operon represses *dnaA* and *dnaN* in *B. subtilis*. This RNA, named ArrA for <u>a</u>nti-sense <u>R</u>NA <u>r</u>epressor of *dna<u>A</u>*, overlaps with the 253 bp 5’ untranslated leader region of the *dnaA* open reading frame and was previously identified in transcriptome analyses and called S2 (Nicolas *et al*., 2012). We found that an *arrA* promoter mutant that had reduced or eliminated production of ArrA RNA caused an increase in DnaA and DnaN mRNA and protein and a reduction in sporulation initiation. We also found that the inhibition of sporulation was alleviated by loss of *sda*, indicating that the effects of *arrA* on sporulation were mediated by its effect on DnaA and subsequent effect on expression of *sda* (Fig 1B). Together, our results indicate that the normal function of *arrA* is to reduce the amount of DnaA and DnaN in the cell and help enable proper initiation of sporulation, and that this antisense control of *dnaA* by *arrA* is critical for normal cellular physiology. Lastly, although there are not many analyses of the small RNAs in various *Bacilli*, ArrA appears to be conserved in at least some members of the *Bacillus* genus.

## Results

### In vitro transcripts from the promoter region of the *dnaA* operon

The promoter region of the *dnaA-dnaN* operon is part of *oriC* and contains several DnaA binding sites (Fig 1A). When DnaA is bound to some of these sites, transcription of the operon is decreased {reviewed in (Kaguni, 2006; Menikpurage *et al*., 2021)}. In the course of analyzing transcription from the *dnaA* promoter (P*dnaA*) in vitro and the activity of DnaA as a transcriptional repressor, we observed a transcript that was more abundant than that initiating from the *dnaA* promoter (Fig 2). The in vitro transcription reactions contained RNA polymerase that had been purified from *B. subtilis*, the major sigma factor SigA, ribonucleotides, appropriate buffer and ^32^P-UTP for visualization of the transcripts (Methods). We used a linear DNA template (a 1,431 bp PCR product) that was expected to give a run-off transcript of ∼250 nucleotides from P*dnaA* (Fig 2A).

**Figure 2:**
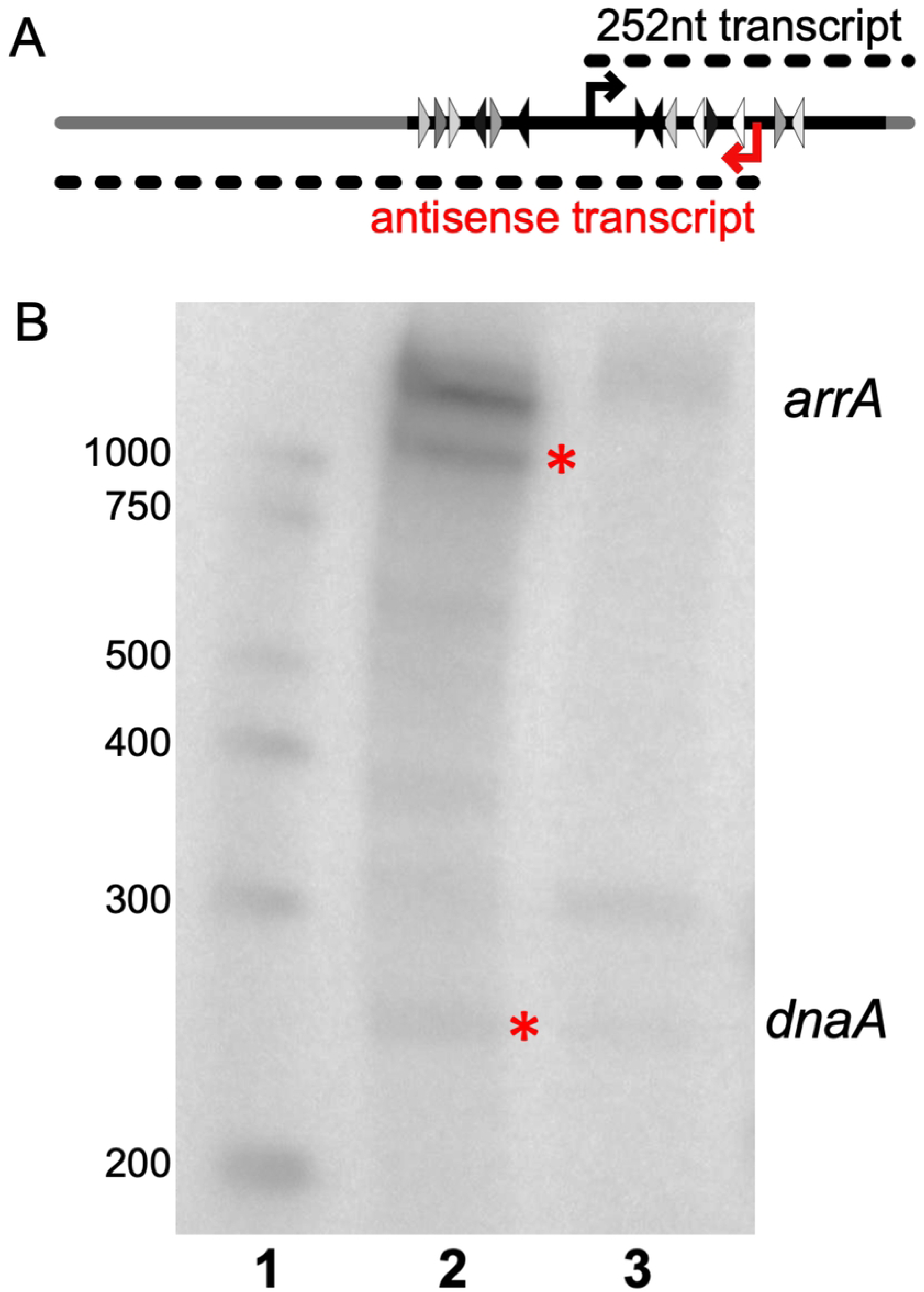
In vitro transcription from the *dnaA* promoter and the antisense *arrA* promoter. **A. Cartoon of the template and transcripts for in vitro transcription.** The linear DNA template used for run-off in vitro transcription was a 1,431 linear PCR product amplified from a plasmid containing the *B. subtilis dnaA* promoter region (enlarged in Fig 1A). This template fragment contains plasmid sequences (gray line) and sequences from the *dnaA* region (black line). Triangles indicate DnaA binding sites as described in Fig 1A. The 252-nucleotide transcript expected from P*dnaA* is depicted as a dotted black line above. The transcript observed that led to the identification of *arrA* is depicted as a dotted line below and labeled in red as antisense transcript. **B. In vitro transcription reactions and repression of P*dnaA* and P*arrA* by DnaA.** In vitro transcription reactions contained 3.5 nM of the DNA template (Fig 2A), 50 nM RNA polymerase and 300 nM sigma-A. Lane 1: Markers, with sizes indicated to the left. Lane 2: Products of in vitro transcription in the absence of DnaA. Products from P*dnaA* and P*arrA* are marked with red asterisks and indicated to the right of the figure. Lane 3: Products of in vitro transcription in the presence of DnaA-ATP (250 nM).

As expected, the in vitro transcription reactions produced a product that was ∼250 nucleotides (Fig 2B, lane 2), corresponding to the predicted product from P*dnaA*. However, we also observed two transcripts (labeled *arrA*) that were >1,000 nucleotides and apparently more abundant than that from P*dnaA*. Based on size, these larger transcripts might initiate from far upstream of P*dnaA* and be in the same orientation, or initiate downstream of P*dnaA* and go in the opposite orientation.

We found that production of the larger transcripts was repressed by DnaA. Addition of DnaA to the in vitro transcription reactions caused a large decrease in the amount of the larger transcripts, and also appeared to modestly repress the transcript from P*dnaA* (Fig 2B lane 3). Repression of both of the larger transcripts by DnaA indicated that they likely initiate from a promoter that is close to DnaA binding sites, making it likely that the promoter was downstream from and in the opposite orientation of P*dnaA* (Fig 2A). We do not know why there are two bands, but they could be isoforms of the same transcript, they could initiate at the same promoter but end at different places, or one could result from end-to-end transcription. Experiments described below allowed us to focus on a single promoter driving a transcript that is anti-sense to the *dnaA* mRNA. Based on results described below, we named the antisense gene *arrA* (pronounced R-A) for <u>a</u>ntisense <u>R</u>NA <u>r</u>epressor of *dna<u>A</u>*.

### The *arrA* promoter

We scanned the DNA sequence downstream from and in the opposite orientation of P*dnaA* for possible promoters that could generate the observed transcript in vitro and be repressed by DnaA. We found a good candidate for a promoter that would likely be recognized by the major form of RNA polymerase (containing SigA). There is a perfect match (5’-TATAAT) to the consensus −10 region and a reasonable match (5’-TacACA) to the consensus −35 region (consensus: TTGACA) with 16 bp separating them (Fig 3A). Further, there are two DnaA binding sites that overlap this promoter, one in −35 region and another between the −10 and −35 regions (Fig 3A), and additional DnaA binding sites downstream from this putative promoter (Fig 2A, 3B).

**Figure 3.**
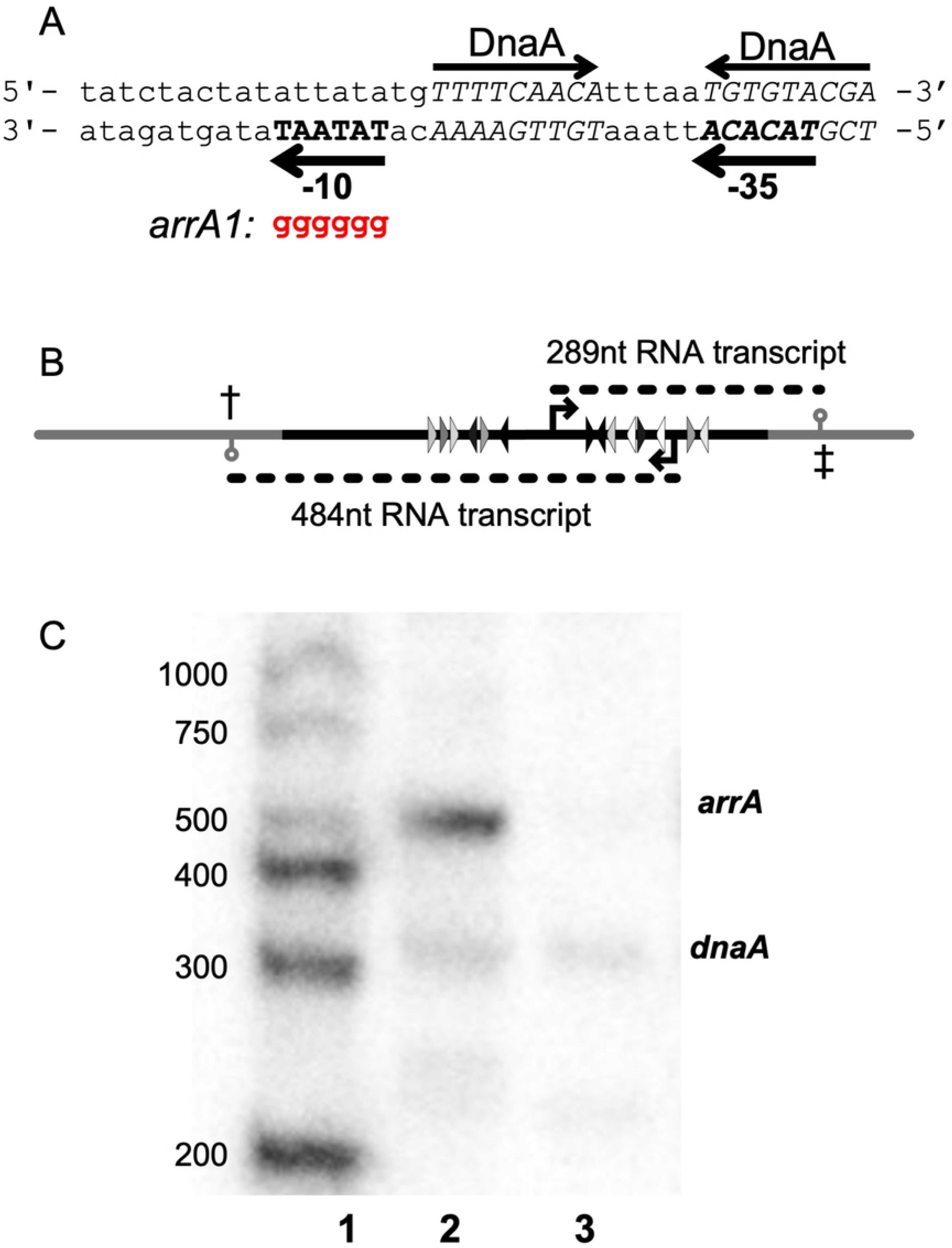
In vitro transcription from the *arrA* promoter. **A. Sequence of the *arrA* promoter region.** The putative −10 and −35 regions of the *arrA* promoter are indicated and shown in bold capitals on the bottom strand going leftward. The −310 sequence is a perfect match to the consensus, and the −35 sequence (TACACA) has two mismatches from the consensus (TTGACA). DnaA binding sites are indicated above the top strand and their sequences shown in italic capital letters. The *arrA1* allele (shown in red) changes the −10 (TATAAT) to GGGGGG. **B. Cartoon of template and transcripts for in vitro transcription.** The template used for in vitro transcription was a 948 linear PCR product amplified from a plasmid containing the *B. subtilis dnaA* and *arrA* promoter regions. This plasmid was engineered to include transcription terminators (⚲): One from *B. subtilis serO/aroC* (‡) downstream from the *dnaA* promoter and another from *gatB/yerO* (†) downstream from the *arrA* promoter (Methods). The portion of the template derived from the *dnaA/arrA* region is shown in black, and the regions derived from the plasmid and the transcription terminators are shown in grey. Other features are as described in Fig 1A. **C. In vitro transcription with wild type and *arrA1* mutant templates.** In vitro transcription reactions contained 2 nM of the DNA template (Fig 3B), 30 nM RNA polymerase and 45 nM sigma-A. Lane 1: Markers, with sizes indicted on the left. Lane 2: Products of in vitro transcription using a template with wild type promoters; transcriptsfrom P*dnaA* and P*arrA* are indicated to the right of the figure. Lane 3: Product of in vitro transcription using a template with the *arrA1* mutant promoter (and wild type *dnaA* promoter).

We found that the putative promoter was responsible for the *arrA* transcript. We altered the −10 region, changing the sequence from 5’-TATAAT to 5’-GGGGGG {*arrA1*} and then used a template containing this mutant promoter in in vitro transcription reactions. In these experiments, we used a 948 bp template that contains both P*dnaA* and P*arrA*, along with transcription terminators that we introduced (Fig 3B; Methods). As expected, there were transcripts of ∼300 and ∼500 nucleotides corresponding to those predicted from P*dnaA* and P*arrA*, respectively (Fig 3C, lane 2). We found that the *arrA1* mutation abolished the *arrA* transcript but had little or no detectable effect on transcription from P*dnaA* (Fig 3C, lane 3). Together, these results indicate that, at least in vitro, there is an antisense RNA that is complimentary to the 5’ untranslated region of the *dnaA* transcript and that this transcript initiates at P*arrA* and is repressed by DnaA.

### *arrA* RNA is made in vivo

The *arrA* transcript has been observed in whole-genome transcriptome studies of *B. subtilis* and was annotated as transcript S2 (Nicolas *et al*., 2012). There is very little ribosome footprint density in this region (Lalanne *et al*., 2018), indicating that the RNA is likely non-coding. In contrast to our observations from in vitro transcription where the *arrA* transcript was more abundant than that from the *dnaA* promoter, there is approximately 20-fold more *dnaA* mRNA than *arrA* RNA during exponential growth in LB and minimal media (Nicolas *et al*., 2012; Lalanne *et al*., 2018). We suspect that the relatively low level of *arrA* RNA in vivo compared to in vitro is likely due to repression of transcription by DnaA in vivo, similar to the relative repression of the two promoters that we observed in vitro (Fig 2B).

We found that production of the *arrA* transcript in vivo was dependent on the sequences we identified as the putative *arrA* promoter. We introduced the *arrA1* allele into its normal chromosomal location in the *dnaA-dnaN* operon (Methods), extracted RNA from cells growing exponentially in LB medium and used Northern blotting to detect the *arrA* RNA (Methods). Transcripts were probed with a radiolabeled oligonucleotide (Fig 4A, indicated as ssDNA oligo) complementary to the 5’ portion of the *arrA* transcript.

**Figure 4.**
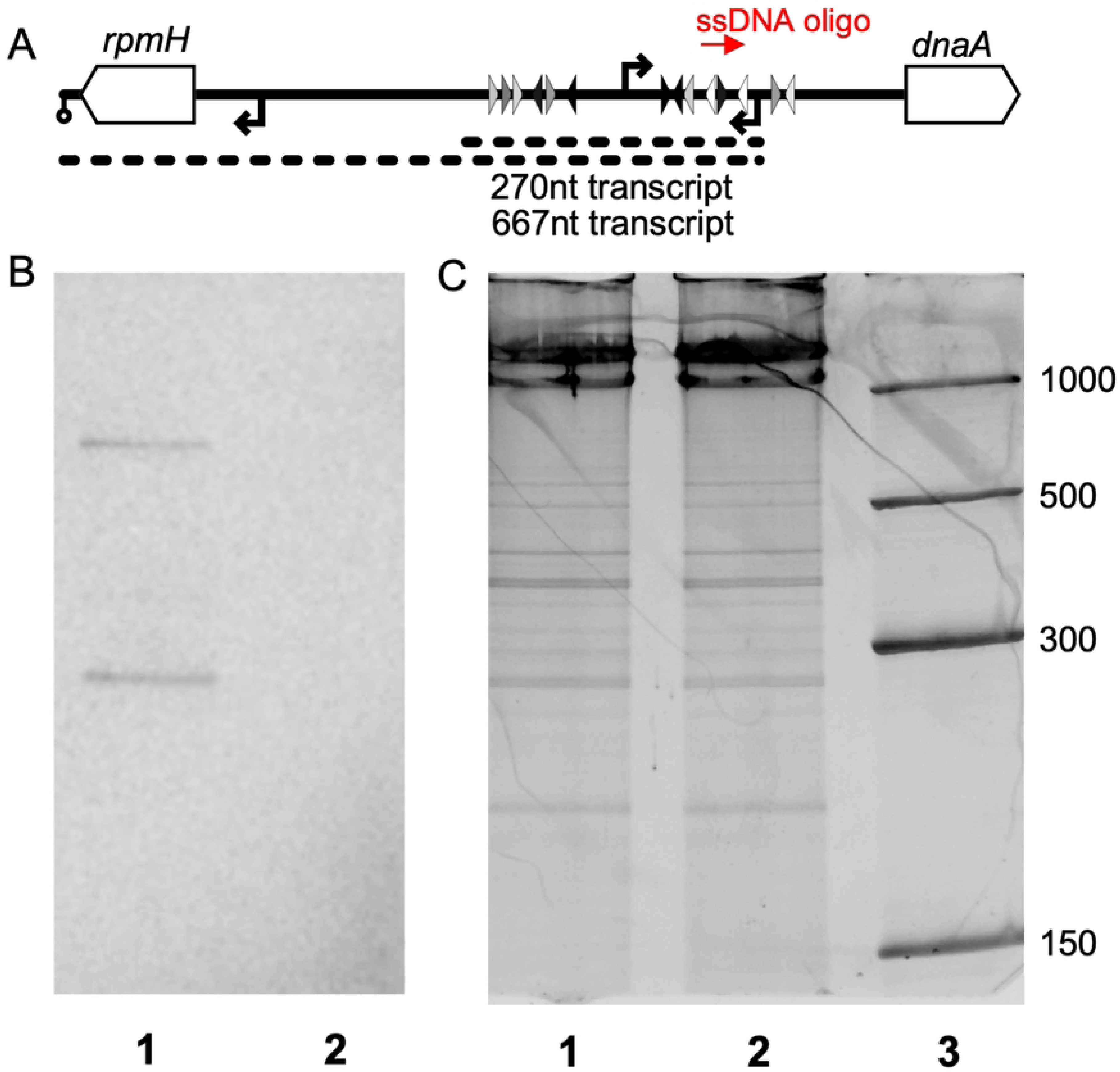
Mutations in the *arrA* promoter region abolish transcription in vivo. **A. Map of the *dnaA* promoter region with probe and transcripts indicated.** *rpmH* and *dnaA* are shown in white (not to scale). The radiolabeled oligonucleotide that was used in Northern blotting hybridizes near the 5’ end of the *arrA* transcript and is indicated in red (ssDNA oligo). A cartoon of the observed transcripts is shown below. The 667 nucleotide transcript extends through *rpmH* and the 270-nucleotide transcript is consistent with either a weak transcriptional terminator or RNA processing of the longer transcript. Promoters and DnaA binding sites are indicated as in figures above. **B. Northern blot probing for *arrA* RNA.** RNA from cells in mid-exponential growth in LB was probed using the radiolabeled ssDNA oligonucleotide that hybridizes near the 5’ end of the *arrA* transcript. Lane 1: RNA from wild type cells. Lane 2: RNA from *arrA1* mutant cells. The small and large bands correspond to the 270 and 667 nucleotide transcripts drawn in panel A. **C.** The same gel in B was stained with SYBR Gold prior to transfer to visualize total RNA. Lane 3 contains RNA markers (sizes in nucleotides on the right). Images of the blot and stained gel are shown in S1A Raw Images.

We observed an approximately 270 nucleotide transcript (Fig 4B lane 1) that hybridized to the *arrA* probe, consistent with transcription initiating at P*arrA* and terminating in the intergenic region between *dnaA* and *rpmH* (Fig 4A). We also observed an ∼670 nucleotide transcript (Fig 4B) consistent with transcription initiating at P*arrA* and extending through *rpmH* and terminating at the *rpmH* terminator (Fig 4A). Notably, both of these transcripts were eliminated in the *arrA1* mutant. That is, there was no detectable RNA corresponding to the two transcripts in this mutant (Fig. 4B, lane 2). There were similar amounts of total RNA in each of the two samples (lanes 1, 2) as determined by staining the gel with SYBR Gold prior to transfer (Fig 4C, lanes 1, 2). We conclude that the *arrA1* mutation reduced transcription of *arrA* to levels below our limit of detection, both in vivo and in vitro, and that the residues changed are most likely the −10 region of the *arrA* promoter.

### Effects of *arrA* on *dnaA* and *dnaN*

We found that *arrA* was important in vivo for proper control of *dnaA* and *dnaN*. We measured the amount of *dnaA* and *dnaN* mRNA in *arrA*^+^ (wild type) and *arrA1* mutant cells during exponential growth in defined minimal medium using RT-qPCR with primers internal to and near the 3’ end of each open reading frame (Methods; S1 Table). We found that cells with the *arrA1* mutation had approximately twofold more mRNA for both *dnaA* and *dnaN* than wild type cells (Fig. 5A). These results indicate that *arrA* normally functions to reduce mRNA levels of *dnaA* and *dnaN*.

**Figure 5.**
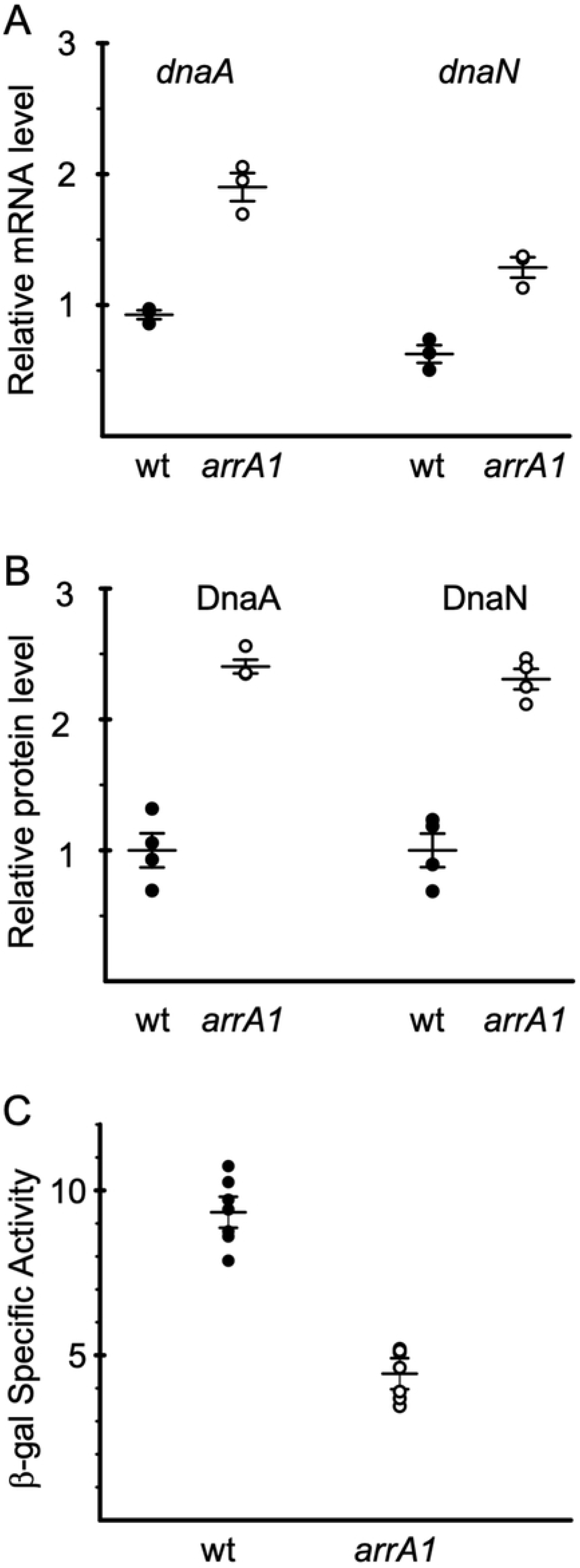
*arrA* affects expression of *dnaA*, *dnaN*, and *ywlC*. Expression of *dnaA* and *dnaN* (mRNA and protein) *ywlC-lacZ* expression were measured in wild type (filled circles) and *arrA1* (open circles) mutant cells during exponential growth in defined minimal medium with glucose. **A. The *arrA1* mutant has elevated levels of *dnaA* and *dnaN* mRNA.** The amount of *dnaA* and *dnaN* mRNA was measured using RT-qPCR. The amount of RNA from each gene was normalized to that from *gyrA.* Error bars show the mean and standard error of the mean from three independent experiments. mRNA from both *dnaA* and *dnaN* is significantly elevated in the *arrA1* mutant (ELS393) compared to that in wild type (AG174) (P-values 0.0057 and 0.0021, respectively, in a two-tailed Student’s *t*-test). **B. The *arrA1* mutant has elevated levels of DnaA and DnaN protein.** The amount of DnaA and DnaN protein was measured by Western blotting. The amount of lysate loaded was normalized to the OD600 of the culture at the time of sampling. The hybridization membrane was cut between the expected molecular weights of DnaA and DnaN and quantitative Western blotting was performed using antibodies against each protein. The data were normalized to the wild type samples for each protein. The amount of both DnaA and DnaN is significantly elevated in *arrA1* cells (ELS393) (four biological replicates) compared to that in wild type (AG174) (P-values of 0.044 and 0.018, respectively, in a two-tailed Student’s *t*-test). An image of the blot is shown in S1B Raw Images. **C. Transcription of *ywlC* is reduced in the *arrA1* mutant.** β-galactosidase specific activity from P*ywlC*-*lacZ* was determined at different times during exponential growth. Data are from two independent cultures for each strain and three or four time points for each culture. Error bars represent the standard error of the mean of all seven samples from each strain. Expression from P*ywlC* was lower in the *arrA1* mutant (grey) compared to wild type (black) (P-value 2.3×10^-7^ in a two-tailed Student’s *t*-test).

The *arrA1* mutant also had increased amounts of DnaA and DnaN protein. We measured the amount of DnaA and DnaN in Western blot experiments using antibodies specific to each protein (Methods). We found that there was approximately twofold more DnaA and DnaN in the *arrA1* mutant than in wild type cells (Fig 5B), consistent with the observed change in the amount of mRNA.

We also found that the *arrA1* mutant had changes in expression of *ywlC*, a gene that is repressed by DnaA (Goranov *et al*., 2005; Ishikawa *et al*., 2007; Breier and Grossman, 2009; Hoover *et al*., 2010; Washington *et al*., 2017). We found that expression of a P*ywlC*-*lacZ* fusion was reduced in the *arrA1* mutant (Fig 5C), likely due to the increased amount of DnaA. Together, our results indicate that *arrA* normally reduces production of DnaA and DnaN by causing a decrease in the amount of *dnaA* and *dnaN* mRNA, and that this regulation is important for proper control of gene expression.

### Little or no effect of *arrA* on growth rate and replication initiation

#### Growth

The growth rate of the *arrA1* mutant was virtually indistinguishable from that of isogenic *arrA*+ cells (Table 1). In rich medium (LB), the doubling times were 21.8 ± 1.3 and 22.3 ±1.3 minutes for wild type and the *arrA1* mutant, respectively. In sporulation medium (DSM), the doubling times were 36 ± 1 and 36.5 ± 0.6 minutes for wild type and *arrA1*, respectively (Table 1; Fig 6A). Similarly, in defined minimal medium with glucose as carbon source (S7), the doubling times were 64 ± 5 for both wild type and the *arrA1* mutant (Table 1). Based on these results, we conclude that there is little to no detectable effect of *arrA* on growth rate under the conditions tested.

**Figure 6.**
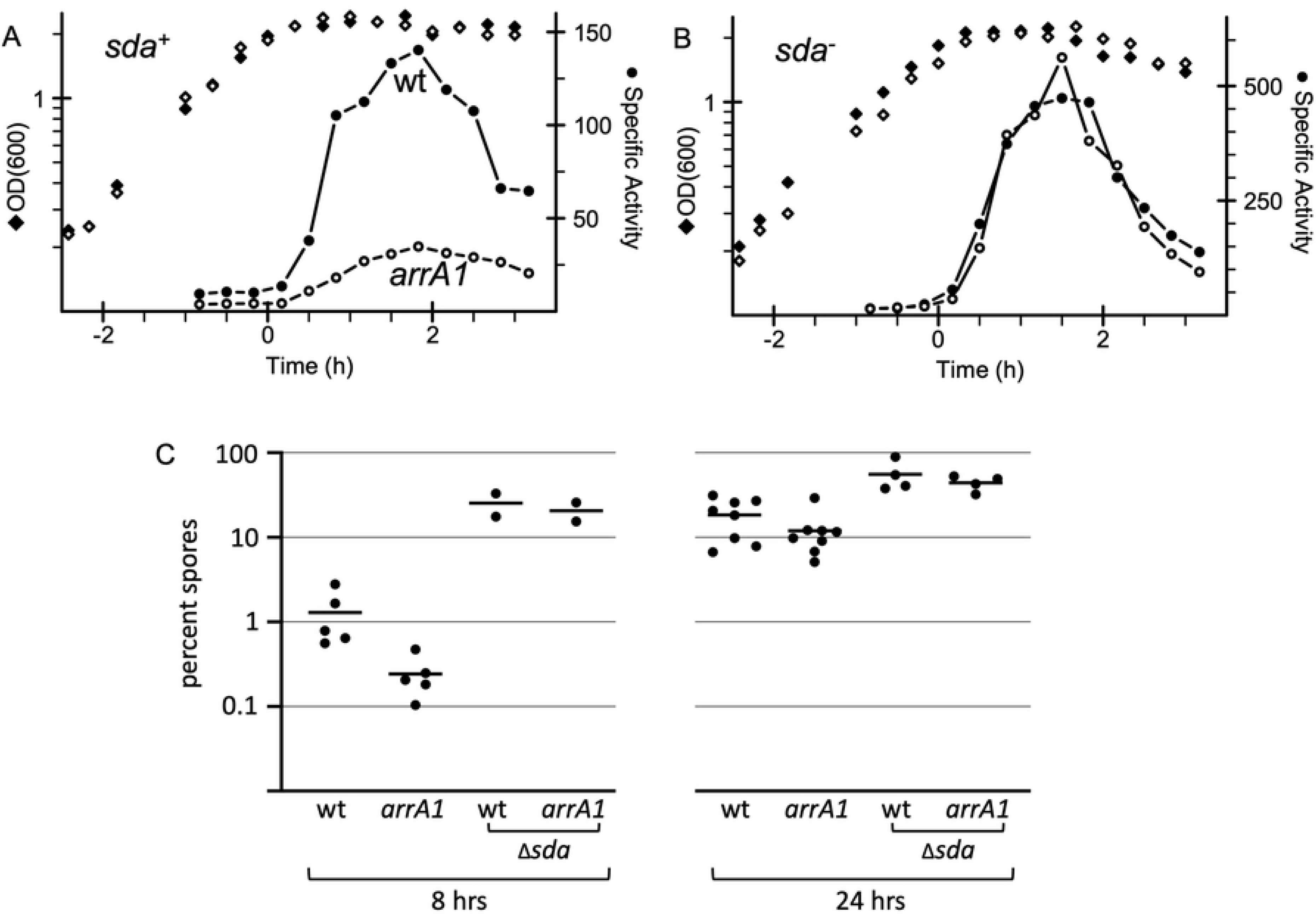
Effects of *arrA* and *sda* on sporulation and early sporulation gene expression. Sporulation and expression of an early sporulation gene (P*spoIIA*-*lacZ*) were measured in cells growing in the sporulation medium DSM. Time 0 represents entry into stationary phase. **A, B.** Growth (OD600; diamonds) and expression of P*spoIIA*-*lacZ* (specific activity; circles) in *arrA*+ (filled symbols) and *arrA1* mutant cells (open symbols). **A.** Wild type *sda* (*sda*^+^) strains: *arrA*^+^ (wt; strain ELS488) and *arrA1* (strain ELS490). One representative experiment of three independent experiments is shown. **B.** Δ*sda* strains: *arrA*^+^ (ELS522), and *arrA1* (ELS526). One representative experiment of two independent experiments is shown. **C.** Percent sporulation (% spores per viable CFU) was determined 8 hrs (left) and 24 hrs (right) after entry into stationary phase. Strains used were: wt (AG174); *arrA1* (ELS393) and Δ*sda arrA*^+^ (ELS484); Δ*sda arrA1* (ELS501). The difference in sporulation between wild type (1.3%) and *arrA1* (0.24%) (both *sda*+) at 8 hrs was statistically significant (P = 0.021 using ratio paired t-tests).

**Table 1.**
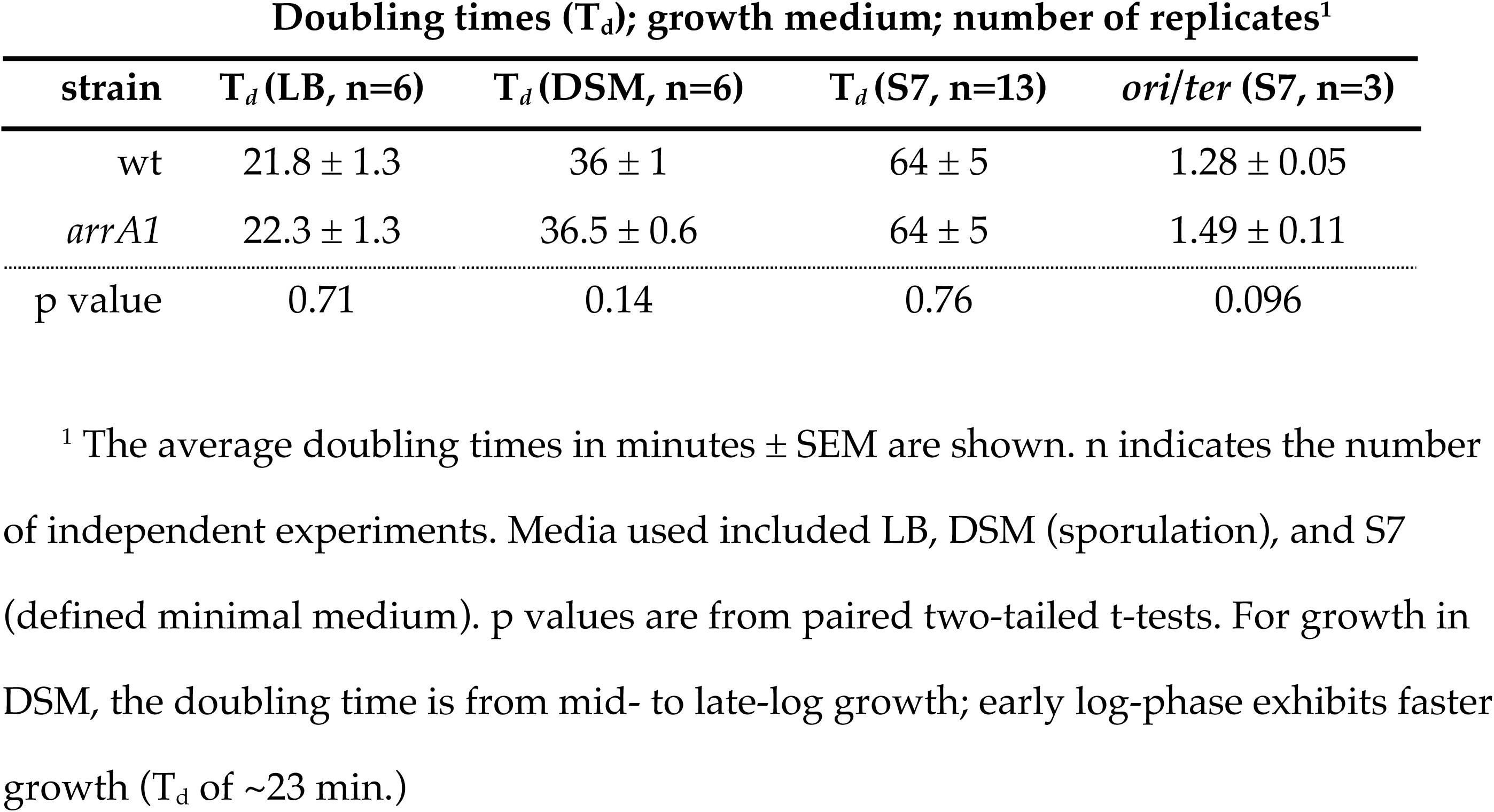
Growth of and replication in *arrA1* mutants.

#### Replication

Increases in the amount of DnaA or DnaN in the cell can cause an increase in replication initiation (Ogura *et al*., 2001; Goranov *et al*., 2009). Therefore, we decided to test if the loss of the *arrA* transcript and subsequent increase in DnaA and DnaN had an effect on replication initiation. We measured the ratio of the amount of DNA from the origin region (*ori*) relative to that of the terminus region (*ter*) by quantitative PCR (qPCR). The ratio of these two regions is a measure of replication and alterations in replication initiation can cause changes in the *ori* to *ter* ratio (*ori*/*ter*). The *ori* to *ter* ratio in cells growing exponentially in defined minimal medium was 1.28 ± 0.05 and 1.49 ± 0.11 for wild type and the *arrA1* mutant, respectively (Table 1). This difference did not appear to be significant (p = 0.096; paired t-test). Based on these results, we conclude that if the *arrA1* mutation causes a change in replication initiation, then this change is within the error of our measurements and more sensitive assays are needed. We have not pursued these. Rather, we focused on the more noticeable effects on gene expression and sporulation.

### *arrA* is needed for normal sporulation initiation

Alterations in the amount of active DnaA (Moriya *et al*., 1990; Lemon *et al*., 2000; Burkholder *et al*., 2001) are known to inhibit sporulation initiation (Ireton and Grossman, 1994; Lemon *et al*., 2000; Burkholder *et al*., 2001). Because *arrA* affects DnaA, we measured its effects on sporulation initiation. We measured expression of *spoIIA*, an operon essential for and expressed early during sporulation, using a fusion of the *spoIIA* promoter to *lacZ* (*spoIIA-lacZ*). We previously used *spoIIA*-*lacZ* fusions to monitor effects of DnaA and replication stress on sporulation (Ireton and Grossman, 1992, 1994; Lemon *et al*., 2000). Transcription of *spoIIA* is activated by the phosphorylated form of the transcription factor Spo0A {reviewed in: (Phillips and Strauch, 2002; Piggot and Hilbert, 2004)} and its expression is reduced under conditions of increased DnaA protein or activity (Lemon *et al*., 2000; Goranov *et al*., 2005; Washington *et al*., 2017).

As expected, in wild type cells transcription of *spoIIA-lacZ* increased shortly after entry into stationary phase under sporulation conditions (Fig. 6A). In contrast, expression of *spoIIA-lacZ* was reduced in the *arrA1* mutant (Fig 6A). These results indicate that *arrA* is important for normal expression of an operon that is essential for and activated early during sporulation.

We also found that production of spores was altered in the *arrA1* mutant. We measured sporulation approximately eight hours after cells entered stationary phase (the start of sporulation). There was an approximately five-fold difference in sporulation between wild type (1.1 x 10^7^ spores/ml; 1.3% sporulation) and the *arrA1* mutant (2.3 x 10^6^ spores/ml; 0.24% sporulation) (Fig 6C). By 24 hours after the initiation of sporulation, there was no significant difference in sporulation levels of wild type (6.5 x 10^7^ spores/ml; 18% sporulation) and the *arrA1* mutant (3.0 x 10^7^ spores/ml; 12% sporulation) (Fig 6C). Based on these results, we conclude that the initiation of sporulation is reduced in the *arrA1* mutant, but that the mutant seems to eventually achieve wild type levels of sporulation, and that the normal function of *arrA* is to enable proper and robust sporulation initiation.

### *arrA* affects sporulation through the sporulation inhibitory gene *sda*

Several mutations and conditions that affect replication and cause an increase in the amount or activity of DnaA also inhibit the initiation of sporulation (Moriya *et al*., 1990; Ireton and Grossman, 1992, 1992; Lemon *et al*., 2000; Burkholder *et al*., 2001; Goranov *et al*., 2005; Washington *et al*., 2017). In these cases, DnaA-mediated inhibition of sporulation is due to increased expression of the sporulation-inhibitory gene *sda*. Transcription of *sda* is directly activated by DnaA (Burkholder *et al*., 2001; Ishikawa *et al*., 2007; Hoover *et al*., 2010) and *sda* encodes a small protein that binds to and inhibits histidine protein kinases needed for activation (phosphorylation) of the transcription factor Spo0A (Burkholder *et al*., 2001; Rowland *et al*., 2004; Whitten *et al*., 2007; Cunningham and Burkholder, 2009) (Fig. 1B).

We found that the effect of *arrA1* on sporulation was mediated by *sda*. In an *sda* null mutant, expression of *spoIIA-lacZ* was virtually indistinguishable between *arrA*^+^ and *arrA1* mutant cells (Fig 6B). Additionally, the sporulation efficiencies of *arrA*^+^ and the *arrA1* mutant were also very similar in the absence of *sda* at both 8 and 24 hours (Fig 6C). Together, our findings indicate that the normal function of *arrA* is to dampen expression of *dnaA*, which in turn leads to decreased expression of the sporulation inhibitor *sda* and enables proper sporulation initiation.

### Possible conservation of *arrA* in some *Bacillus* species

We searched the literature for evidence of possible small antisense RNA in the *dnaA* promoter region in other organisms. Many transcriptome analyses have not captured small RNAs, but of those that did, ArrA-like RNAs were not detected in *Escherichia. coli*, *Vibrio natrigens*, *Caulobacter crescentus* (Lalanne *et al*., 2018), *Staphylococcus. aureus*, members of the *Clostridia* genus, several different strains of *Streptococcus pneumoniae*, or *Enterococcus faecalis* (Pichon and Felden, 2005; Mraheil *et al*., 2010; Tsui *et al*., 2010; Shioya *et al*., 2011; Acebo *et al*., 2012; Mann *et al*., 2012, 2012; Soutourina *et al*., 2013; Bronsard *et al*., 2017; Sinel *et al*., 2017; Sinha *et al*., 2019; Subramanian *et al*., 2019).

#### Antisense RNA in the *dnaA* region in *B. licheniformis*

An antisense RNA in the *dnaA* region was identified in *B. licheniformis* in transcriptome analyses (Wiegand *et al*., 2013; Guo *et al*., 2015). Additionally, we identified a potential promoter for the antisense RNA between *dnaA* and its promoter (**Fig 7A)**. There is a possible −10 region (5’-TATAGT) that has one mismatch from consensus (5’-TATAAT) and a possible −35 region (5’-ATGAAA) that has two mismatches from consensus (5’-TTGACA). Furthermore, there are two DnaA binding sites flanking the putative −10 region that are identical to those in *B. subtilis* (Fig 7A). Overall there is high sequence similarity in this region between *B. subtilis* and *B. licheniformis*, including the spacing of the two DnaA binding sites, although the two predicted promoters are offset by 5 bp. Based on this information, we suspect that *B. licheniformis* contains *arrA* and it is regulated and functions similarly to that of *B. subtilis*.

**Figure 7.**
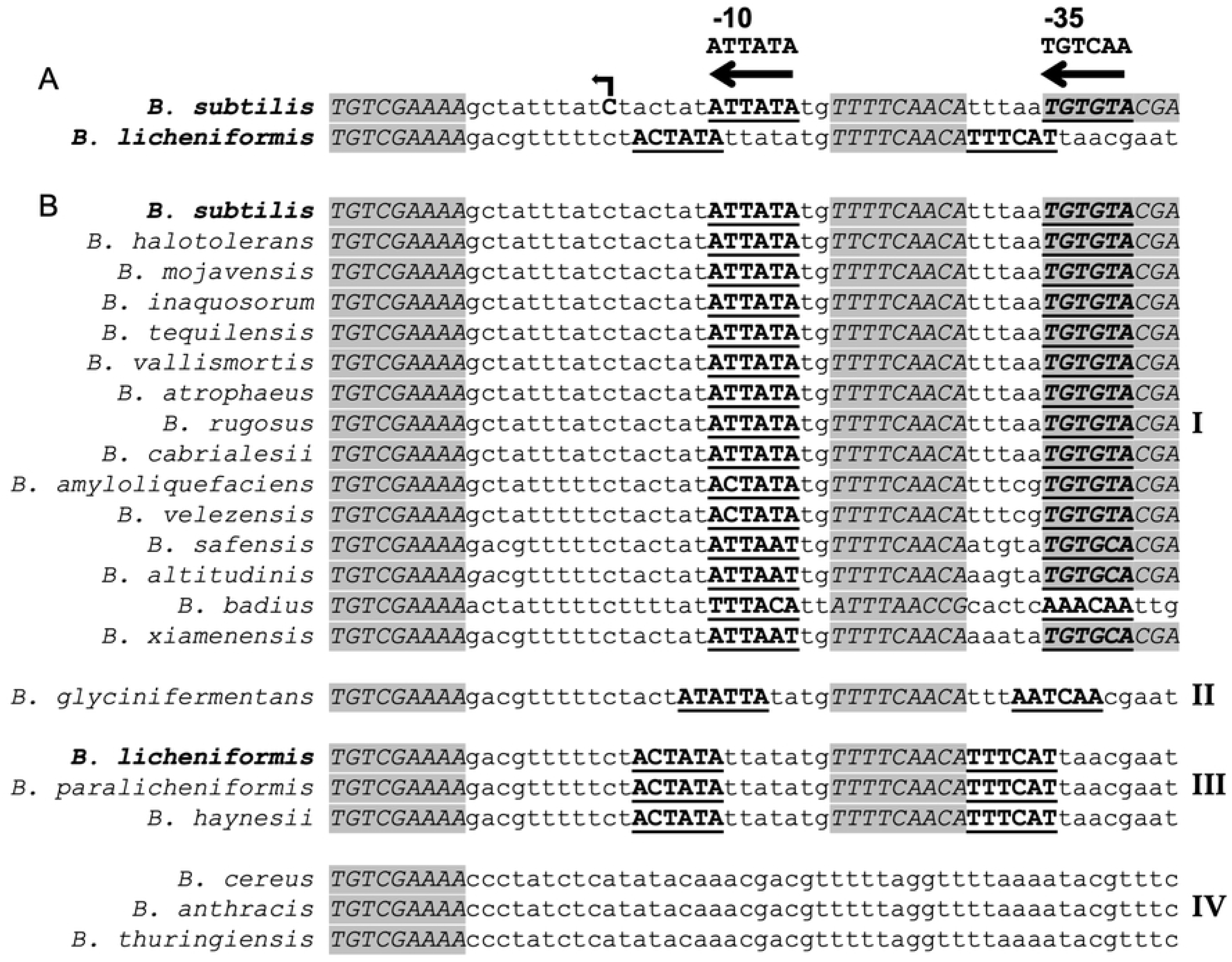
Identification of possible antisense promoters with DnaA binding sites in various *Bacillus* species. Sequences (5’ to 3’) from downstream of the *dnaA* promoter and in the vicinity of the *arrA* promoter from *B. subtilis* were compared and aligned using the ClustalOmega algorithm in JalView. We aligned the sequences by the conserved DnaA binding site (at the left of each sequence shown). Sequences matching a DnaA binding site are in capital letters, italicized, and shaded grey. The sequences of putative promoters (−10 and −35) for an antisense RNA (*arrA*) are shown in capital letters, bold and underlined. The consensus promoter sequence is shown above. Note: these are drawn on the anti-sense strand to *dnaA*, so the sequences are complimentary to the traditional −10 (TATAAT) and −35 (TTGACA). **A.** Comparison of the regions from *B. subtilis* and *B. licheniformis*. The *B. subtilis arrA* transcription start site is in bold, with a bent arrow on top indicating the direction of transcription. **B.** Sequences from different species are divided into four groups based on similarities and differences to *B. subtilis* and *B. licheniformis*. Groups I, II, and III, have two or three DnaA binding sites and the spacing between them is conserved. They all have a putative promoter that is antisense to *dnaA* and in all cases, the spacing between the −10 and −35 is 16 bp. **I.** Regions from 14 *Bacillus* species that are most similar to that in *B. subtilis.* There are 16 bp between the putative −10 and the downstream DnaA binding site, and 2 bp between the −10 and the upstream DnaA binding site. **II.** There is one member of this group, and the sequences are very similar to Group I, but the putative promoter elements are shifted by 2 bp leaving 14 bp between −10 and the downstream DnaA binding site and 4 bp between the −10 and the upstream DnaA binding site. **III.** Regions from two species are most similar to that from *B. licheniformis*. There are 11 bp separating the putative −10 and the downstream DnaA binding site, and 7 bp separating the putative −10 and the upstream DnaA binding site. **IV.** The *arrA* promoter does not appear to be present in *B. cereus* and members of its phylogenetic group, including *B. anthracis* and *B. thuringiensis*.

#### Possible antisense promoter in the *dnaA* region of other *Bacilli*

We analyzed the region between the promoter and *dnaA* of 36 different *Bacillus* species (Methods; S2 Table) for possible promoters and DnaA binding sites, similar to those found in *B. subtilis* and *B. licheniformis* (Fig 7A). We found sequences similar to the consensus promoter in 17 additional regions analyzed (of 36) and grouped these based on whether the spacing between the −10 and the downstream DnaA binding site was more like that of *B. subtilis* (16 bp) (Fig 7B, Group I) or *B. licheniformis* (11 bp) (Fig 7C). Fourteen had 16 bp (Fig. 7B, Group I) and one had 14 bp (Fig 7B, Group II) between the putative −10 and the downstream DnaA binding site identical to that in *B. subtilis*. Two organisms had spacing between the putative −10 and the downstream DnaA binding site that is identical to that in *B. licheniformis* (Fig 7B, Group III). In all cases, the distance between the upstream and downstream nucleotide of the two DnaA binding sites (one downstream of the −10 and the other between the −10 and −35) is 24 bp even though the exact spacing relative to the putative promoter is somewhat different. We did not find similar promoter-like sequences for a possible antisense RNA in the *dnaA* region of the *Bacillus cereus* group, including *B. cereus, B. anthracis*, and *B. thuringiensis* (Fig 7B, Group IV). Together, these analyses indicate that an *arrA*-like transcript is likely made in several, but not all, *Bacillus* species and that its expression is repressed by DnaA. We suspect that its function in the modulation of expression of the *dnaA-dnaN* operon is conserved.

## Discussion

We identified *arrA* as an antisense regulator of the *dnaA* operon. Transcription of *arrA* is repressed by DnaA and loss of *arrA* transcription caused an increase in the amount of DnaA and DnaN and a decrease in sporulation initiation. Below, we discuss *arrA* in the context of mechanisms regulating *dnaA*, sRNA, and its conservation and effects on sporulation.

### Strategies for regulating DnaA

Bacteria use many different mechanisms to regulate DnaA levels and activity. Different organisms use different combinations; some mechanisms are highly conserved, whereas others are specific to particular taxa. For example, autoregulation by DnaA to repress its own promoter is highly conserved (Menikpurage *et al*., 2021). Also, in all organisms, the ATP-bound form of DnaA is needed for cooperative and proper binding to *oriC* to enable replication initiation {reviewed in: (Mott and Berger, 2007; Chodavarapu and Kaguni, 2016; Rashid and Berger, 2025). Although virtually all organisms regulate cooperative binding of DnaA to *oriC*, the mechanisms used to control binding can be quite different (Jameson and Wilkinson, 2017; Felletti *et al*., 2019). For example, *E. coli* and related species regulate ATP hydrolysis and exchange, have mechanisms to titrate DnaA from *oriC*, and can sequester *oriC* to inhibit access by DnaA. In contrast, *B. subtilis* and apparently many other Gram positive bacteria use anti-cooperativity proteins to modulate proper binding of DnaA to *oriC*. Additional mechanisms modulating DnaA include regulated proteolysis (Gorbatyuk and Marczynski, 2005) and trans-translation (Cheng and Keiler, 2009).

### Small RNAs and *dnaA*

Small RNAs have emerged as a ubiquitous method of gene regulation, especially for the purposes of fine-tuning expression and in stress responses (Adams and Storz, 2020; Jørgensen *et al*., 2020; Papenfort and Melamed, 2023). sRNAs have been identified in *dnaA* and *dnaN* in a few organisms. For example, there is an antisense RNA with unknown function internal to *dnaN* in *E. coli* (Lalanne *et al*., 2018). An antisense RNA produced from within *dnaA* in *Salmonella enterica* appears to increase production of DnaA, perhaps by stabilizing its mRNA (Dadzie *et al*., 2013). There are two sRNAs in *C. crescentus* that are predicted to work in trans and target *dnaA* (Beroual *et al*., 2018), and a trans-acting sRNA regulates both *dnaA* and the master cell cycle regulator *gcrA* in members of the genus *Sinorhizobium* (Jiménez-Zurdo and Robledo, 2015; Robledo *et al*., 2015). These examples point to the utility of sRNAs as a method of replication and cell cycle control.

### DnaA and sporulation in *B. subtilis*

Alterations in DnaA activity have profound effects on cellular physiology and development. In addition to acting as the replication initiator, DnaA acts as a transcription factor and in this way, plays an important role in regulating the initiation of sporulation in *B. subtilis* (and likely other endospore-forming bacteria).

Entry into the sporulation pathway is highly regulated. The key signals to start sporulation include nutrient depletion and high population density (Grossman, 1995; Piggot and Hilbert, 2004). Additionally, conditions that alter replication can inhibit the initiation of sporulation (Ireton and Grossman, 1992, 1994; Lemon *et al*., 2000). These alterations lead to an increase in active DnaA and/or the RecA-dependent host SOS response, both of which cause increased transcription of *sda*, the product of which is an inhibitor of sporulation initiation. The function of *sda* is most easily seen following DNA damage or perturbations to replication, both of which cause an increase in the activity of DnaA (Burkholder *et al*., 2001; Ishikawa *et al*., 2007; Hoover *et al*., 2010). Transcription of *sda* is also controlled in a cell-cycle-dependent manner and very small changes in the amount of DnaA apparently affect transcription of *sda* (Veening *et al*., 2009).

Our results indicate that modest (2-fold) changes in the amount of DnaA and DnaN due to *arrA* affect sporulation initiation. These effects are most likely mediated by DnaA as a direct transcriptional activator of *sda*. However, we suspect that DnaN might also play a role in this. DnaN indirectly affects the activity of DnaA by inhibiting YabA, a protein that normally inhibits cooperative DNA binding by DnaA (Merrikh and Grossman, 2011; Scholefield and Murray, 2013). We suspect that the combined effects of the increase in DnaA and DnaN cause the *sda*-dependent inhibition of sporulation initiation.

*arrA* appears to be conserved in several *Bacillus* species as at least 17 have a candidate promoter and associated DnaA binding sites analogous to those in *B. subtilis*. It is not surprising that *arrA* is not conserved beyond these species. sRNAs appear to be found in most bacterial species where they have been examined. *B. subtilis* makes many different sRNAs (Irnov *et al*., 2010; Dambach *et al*., 2013; Brantl and Brückner, 2014; Lawaetz *et al*., 2024), several of which are antisense to functional genes and known to modulate gene expression (Brantl and Müller, 2021). Despite their prevalence, sRNAs are not frequently conserved, even within members of one genus or between strains within the same species. For example, *B. licheniformis* produces a species-specific sRNA to regulate subtilisin production whereas its close relative *B. subtilis* does not (Hertel *et al*., 2017). Similarly, an RNAseq study of sRNAs in *Rhodobacter capsulatus* found 422 putative sRNAs, of which 298 appear to be species specific and only 40 have putative inter-taxa homologs (Grüll *et al*., 2017). This lack of conservation is thought to reflect the utility of sRNAs as adaptable fine-tuning mechanisms, possibly evolving quickly according to the needs of individual species or strains. Our results highlight the importance of having multiple mechanisms to regulate *dnaA*, the evolutionary diversity of those mechanisms, and the critical role DnaA plays in the cellular decision to enter into the complex process of sporulation.

## Methods

### Media and growth conditions

LB medium was used for routine maintenance of both *E. coli* and *B. subtilis.* Standard concentrations of antibiotics were used for selection when appropriate (ed. CR Harwood, and SM Cutting, 1990). For many experiments (as indicated in the text), *B. subtilis* cells were grown with shaking at 37°C in S7_50_ defined minimal medium with 50 mM MOPS (3-(N-morpholino)propanesulfonic acid) buffer supplemented with 1% glucose, 0.1% glutamate, trace metals, 40 μg/ml phenylalanine, and 40 ug/ml tryptophan (Jaacks *et al*., 1989). *B. subtilis* cultures in minimal medium were typically started from cells grown overnight on solid medium to a very light lawn, resuspended and then used to inoculate liquid cultures at OD600 0.05. Cultures grown in LB medium were inoculated with a single colony from a freshly streaked LB agar plate.

### Sporulation assays

Cells were grown in Difco Sporulation Medium (DSM) (ed. CR Harwood, and SM Cutting, 1990). At the indicated times after entry into stationary phase, samples were diluted and the number of CFUs were determined by plating on LB agar before (total cells) and after (spores) heating to 80°C for 10 min. Percent sporulation was determined by the ratio of spores/ml (CFU/ml after heat treatment) to cells/ml (CFU/ml before heat treatment) x 100%.

### Strains and alleles

*B. subtilis* strains (Table 2) were derived from JH642 (*pheA1 trpC2*) (Perego *et al*., 1988; Smith *et al*., 2014) and were constructed by natural transformation.

**Table 2.** *B. subtilis* strains used.

| Strain | Genotype (reference) |
| --- | --- |
| AG174 | *phe trp* (Perego *et al.*, 1988; Smith *et al.*, 2014) |
| ELS393 | *arrA1 phe trp* |
| MEA448 | *amyE::*(PywlC-lacZ cat) *phe trp* |
| ELS446 | *amyE::*(PywlC-lacZ cat) *arrA1 phe trp* |
| ELS484 | Δ*sda::spc phe trp* |
| ELS488 | SPβ::(*spoIIA-lacZ cat erm*) *phe trp* (Ireton *et al.*, 1994) |
| ELS490 | SPβ::(*spoIIA-lacZ cat erm*) *arrA1, phe trp* |
| ELS501 | *arrA1* Δ*sda::spc phe trp* |
| ELS522 | Δ*sda::spc* SPβ::(*spoIIA-lacZ cat erm*) *phe trp* |
| ELS526 | Δ*sda::spc* SPβ::*spoIIA-lacZ cat erm*) *arrA1 phe trp* |

#### arrA1

*arrA1* alters the −10 region of the *arrA* promoter, changing the 5’-TATAAT to 5’-GGGGGG. It was made by cloning a PCR fragment containing the *arrA* promoter region into the plasmid pCAL1422 (*lacZ, cat*) (Thomas *et al*., 2013), introducing the promoter mutation, and then transforming into *B. subtilis* selecting for chloramphenicol resistance. Transformants have the plasmid integrated by a single crossover and were *lacZ^+^*. A transformant was passaged in LB (without antibiotic) and screened for the loss of *lacZ*, indicating the loss of the integrated plasmid. The *arrA* promoter region was PCR amplified from the chromosome of several isolates that had lost *lacZ* and sequenced to identify a clone that contained the desired mutation in P*arrA*.

#### PywlC-lacZ

A DNA fragment from 240 bp upstream to 73 bp downstream from the start codon of *ywlC* was PCR-amplified and combined with a chloramphenicol resistance cassette, *lacZ*, and upstream and downstream homology regions for insertion into *amyE* using Gibson Isothermal Assembly (Gibson *et al*., 2009). These additional components were PCR-amplified using pDG268 as a template (Antoniewski *et al*., 1990). The linear assembly was then integrated into the *B. subtilis* chromosome at *amyE* by double crossover, selecting for chloramphenicol resistance.

#### Δsda::spc

*Spc* was cloned between sequences flanking *sda* and then a PCR product containing the allele was introduced into the *B. subtilis* chromosome by transformation and selection for spectinomycin resistance. The Δ*sda*::*spc* allele is missing *sda* from 40 bp upstream to 142 bp downstream from the start codon of the 138 bp *sda* open reading frame, essentially as described (Burkholder *et al*., 2001).

#### SPβ::spoIIA-lacZ

*spoIIA-lacZ* was previously described (Wu *et al*., 1991; Ireton and Grossman, 1992). Briefly, it is a transcriptional fusion of the *spoIIA* promoter region fused to *lacZ*, integrated into the temperate phage SPβ.

### Protein expression and purification

#### B. subtilis DnaA

*B. subtilis* DnaA was purified as described previously (Smith and Grossman, 2015).

#### *B. subtilis* RNA polymerase

RNA polymerase was purified from the *B. subtilis* strain MH5636 that expresses *rpoC*-*his*_10_ (Qi and Hulett, 1998), essentially as described (Birch *et al*., 2017), with a few modifications. Briefly, the cell resuspension buffer was supplemented with 5 mM imidazole and β-mercaptoethanol reduced to 2 mM. The extract was clarified using one round of centrifugation followed by filtration through a Whatman FP30/1.2CA filter unit. Talon Superflow resin (Cytiva) was used in place of Ni-NTA, and Heparin and HiQ chromatography was carried out at room temperature, with fractions collected in a 4°C cabinet. The final protein concentration was determined using the Bio-Rad Protein Assay standardized to BSA, and the preparation was stored at −20°C.

#### Sigma-A

*B. subtilis* sigma-A (SigA) protein was isolated from strain JLS116, an *E. coli* strain previously described (Nakano *et al*., 2006) that overexpresses a SigA-chitin fusion protein. Purification was carried out essentially as described (Nakano *et al*., 2006), with minor modifications. Triton X-100 was omitted from the lysis buffer until after lysis by French press was complete. In addition, a protease inhibitor cocktail and 250U of Benzonase nuclease (both from Millipore Sigma) were added immediately after lysis. After centrifugation the lysate was passed through a Whatman FP30/1.2 CA filter unit prior to incubation with the chitin resin. The final preparation was concentrated using a 15-ml Centricon filter (3-kDa cutoff; Amicon), dialyzed against the published storage buffer except that glycerol was increased to 50%, and stored at −20°C. The final protein concentration was determined using the Bio-Rad Protein Assay standardized to BSA.

### Templates for in vitro transcription

Templates for in vitro transcription reactions contained DNA from the *dnaA/arrA* promoter region as indicated in the relevant figures (Figs 2, 3). Template DNA in each in vitro transcription reaction was a linear PCR product of the indicated region that was made using plasmid DNA as a template for PCR using KAPA HiFi HotStart ReadyMix (Roche Sequencing Solutions). PCR products were purified using a Qiagen MinElute PCR cleanup kit and DNA was eluted in nuclease-free water.

Plasmids pELS33, pELS51, and pELS69 were used to generate the PCR products. pELS33 contains the promoter region from 398 bp upstream to 18 bp downstream and pELS51 and pELS69 contain the promoter region from 547 bp upstream to 18bp downstream of the *dnaA* start codon. pELS51 and pELS69 also contain two bidirectional terminators, one at each end of the promoter-containing fragment. Terminators from the *gatB/yerO* (from 2 bp to 34 bp downstream of the *gatB* stop codon) and *serA/aroC* (from 2 bp to 36 bp downstream of the *serA* stop codon) regions were synthesized as oligonucleotides and cloned into each of the two plasmids via Gibson isothermal assembly (Gibson *et al*., 2009). pELS51 and pELS69 are essentially the same, except pELS69 contains the *arrA1* mutant promoter (5’-GGGGGG in place of 5’-TATAAT in the −10 region of the promoter).

### In vitro transcription

In vitro transcription reactions were done in 20 µl reactions that contained 40 mM HEPES-KOH pH 7.6, 10 mM magnesium acetate, 10 mM sodium chloride, 10 mM potassium chloride, 2% sucrose, 2.5% glycerol, 0.5 mM EDTA, 150 mM potassium glutamate, 1 mM DTT, and 0.2 U/µl RnaseOUT. A mix of nucleotides was prepared than then added to transcription reactions at final concentrations of: 2.5 mM ATP; 0.1 mM each CTP, GTP, and UTP, and 1 µC of alpha-^32^P-UTP.

The concentrations of DNA template, RNA polymerase, sigma-A, and DnaA-ATP are indicated in figure legends. RNA polymerase, sigma-A, and RNaseOUT were mixed prior to addition to transcription reactions. Where added, DnaA was pre-incubated on ice for 20 minutes with 2.5 mM ATP in DnaA storage buffer (45 mM HEPES-KOH pH 7.6, 10 mM magnesium acetate, 0.5 mM EDTA, 700 mM potassium glutamate, 1 mM DTT). DnaA-ATP (250 nM) was then incubated on ice for 20 minutes with the template DNA and all remaining components listed above. The RNAP mixture was then added to the final concentrations indicated in the figure legends and incubated for another 20 minutes on ice. The reactions were finally started by addition of the nucleotide mix and the reactions were incubated at 37°C for 20 minutes. Reactions were stopped by adding 10 µl of loading buffer (7 M urea, 100 mM EDTA, 4.8% glycerol, 0.25 mg/ml bromophenol blue, 0.25 mg/ml xylene cyanol) and heating to 75°C for 3 minutes. Samples were run on a polyacrylamide gel containing 7 M urea in tris-borate EDTA running buffer. Gels were dried onto Whatman paper and used to expose a Kodak Storage Phosphor Screen SD230 overnight before imaging on a GE Typhoon FLA 9500 scanner.

RNA size standards were transcribed using Century Marker Template Plus fragments (Invitrogen by ThermoFisher Scientific, AM7782) and an Ambion MAXIscript T7 In Vitro Transcription Kit (AM1312) according to the manufacturer’s instructions.

### β-galactosidase assays

Cells were grown in defined minimal medium at 37°C to mid-exponential phase, diluted back to OD600 0.05, grown to mid-exponential phase (OD 0.2-0.4) and samples were collected. Growth was stopped and cells permeabilized by addition of toluene (1.5% final concentration). β-galactosidase specific activity ({ΔA420 per min per ml of culture per OD600} × 1000) was determined essentially as previously described (Miller, 1972).

### RNA isolation and reverse transcription

Cells were grown to mid-exponential phase overnight on solid medium and used to inoculate liquid cultures at OD600 0.05. When cultures reached OD600 0.2-0.5, 1 OD600 of cells was harvested in an equal volume of ice-cold methanol and pelleted. Pellets were resuspended in 200 µl TE pH 8.0 (10 mM Tris-HCl, 1 mM EDTA) with 10 mg/ml lysozyme and incubated for 5 minutes at 37°C. RNA was then isolated using Qiagen RNeasy PLUS kit. Genomic DNA was removed using dsDNase (Thermo Scientific, EN0771). iScript Supermix (Bio-Rad) was used for reverse transcriptase reactions to generate cDNA. RNA was degraded by adding 75% volume of 0.1 M NaOH and incubating at 70°C 10 minutes, then neutralized by adding 75% of the original volume 0.1 M HCl. RNA was normalized on a Thermo Scientific™ NanoDrop™ One.

### RT-qPCR and qPCR

RT-qPCR and qPCR were done using SsoAdvanced Universal SYBR master mix and CFX96 Touch Real-Time PCR system (Bio-Rad) using indicated primer pairs (S1 Table).

### Northern blots

Total RNA for northern blots was purified (as above) from 2-2.5 OD600 units of cells grown in LB medium to mid-exponential phase. Twenty micrograms of RNA in RNA loading dye (New England Biolabs) per lane was run on a 5% acrylamide (19:1 acrylamide:bisacrylamide ratio) gel containing 8M urea and 1xTBE buffer. Size standards (5 µl of Low Range ssRNA Ladder; New England Biolabs) were also run. Gels were stained with SYBR Gold (Invitrogen) and imaged using a UV transilluminator. RNA was transferred to Hybond-N+ membrane (Cytiva) using a BioRad semidry transfer apparatus and then crosslinked using a UV Stratalinker (Stratagene). The membrane was incubated overnight at 42°C in ULTRAhyb Ultrasensitive Hybridization Buffer (Invitrogen) with an oligonucleotide (oJLS184; 5’-cacagcttgtgtagaaggttgtccacaag) that was end-labeled (T4 polynucleotide kinase, New England Biolabs) with ATP [γ-^32^P] (PerkinElmer). This oligonucleotide is complementary to the 5’ portion of the *arrA* transcript (Fig 4A.) The membrane was washed at 42°C in 2X SSC, 0.5% SDS, exposed to a phosphorstorage screen (GE Life Science) and imaged with a laser scanner (Typhoon FLA9500, GE Life Sciences).

### DNA isolation

Cultures were grown to mid-exponential phase (OD600 0.2-0.4) in defined minimal medium at 37°C. Cells were harvested in one volume of ice-cold methanol, pelleted, and stored at −80°C. Genomic DNA was isolated using Qiagen DNeasy kit with 40 μg/ml lysozyme.

### Western blots

Cultures were grown to mid-exponential phase overnight on solid medium and used to inoculate liquid cultures at OD600 0.05. When cultures reached OD600 0.2-0.5, 1 OD600 of cells was harvested and lysed with 1µg lysozyme and P8849 protease inhibitor cocktail at 37°C. Due to the difficulty of accurately measuring protein concentration following addition of lysozyme, lysates were normalized to culture OD and mixed with ⅓ volume of 4X protein loading buffer (200 mM Tris pH 6.8, 8% SDS, 40% glycerol, 20% β-mercaptoethanol, 0.002% bromophenol blue). Equal volumes of each normalized lysate from four biological replicates were separated on a single 10% polyacrylamide gel and transferred to a nitrocellulose membrane using the Trans-blot SD semi-dry transfer cell (Biorad). The membrane was washed in water and then incubated in Revert 700 Total Protein Stain. After washing with Revert 700 Wash Solution, the stained blot was imaged on a LiCor scanner. After imaging, the blot was washed in water before proceeding with western blotting. According to the manufacturer’s instructions, blots were blocked with ½ X Odyssey Blocking Buffer (diluted 1:1 in phosphate-buffered saline) for 1 hr at room temperature. Blots were then incubated with primary antibody (rabbit 1:10,000 in ½ X Odyssey Blocking Buffer with 0.2% Tween) overnight at 4°C. The primary antibodies were rabbit polyclonal antibodies made against purified DnaA and DnaN (Covance). Blots were washed 5 times with phosphate-buffered saline + 0.2% Tween for at least 5 min each and then incubated for 30 min at room temperature with secondary antibody (LiCor dye 800 goat anti-rabbit 1:10,000 in ½ X Odyssey Blocking Buffer + 0.2% Tween + 0.01% SDS). Blots (S1 Raw Image) were imaged and quantitated on a LiCor scanner.

Quantitation of total protein, DnaA, and DnaN was performed in ImageJ using band densitometry. Each DnaA and DnaN band was normalized to the total protein signal between 250 kD and 20 kD in their respective lanes. DnaA bands were normalized to the average of all wildtype DnaA bands in order to display the fold change between wild type and the *arrA1* mutant. DnaN was normalized similarly.

### Search for a possible antisense promoter and DnaA binding sites between *dnaA* and its promoter

Reference genomes were accessed through the NCBI National Library of Medicine Genome Browser. The search term “Bacillus (Bacillus rRNA group 1)” was entered into the taxonomic search bar and filtered by Status and Assembly Level to show only reference genomes assembled to the chromosome level or beyond. Metadata for 37 “Bacillus” results were downloaded along with RefSeq files (Sequences and Annotations in GBFF format). Reference genomes *B. subtilis* subsp. *subtilis* and *B. subtilis* subsp. *spizizenii* were omitted as they are identical to our wildtype strain, AG174, in the regions compared. We extracted the antisense sequence of the intergenic region between *dnaA* and *rpmH* from each file, aligned them using the ClustalO algorithm in JalView. We then looked for conservation of the DnaA binding sites and scanned the alignment for possible promoters, analogous to those present in *B. subtilis* and *B. licheniformis*.

## Acknowledgments

We thank Mary Anderson for helpful discussions and comments. Peter Zuber and Cierra Birch for strains and advice on purifying RNA polymerase and Sigma-A.

## Supporting information

**S1 Table.**
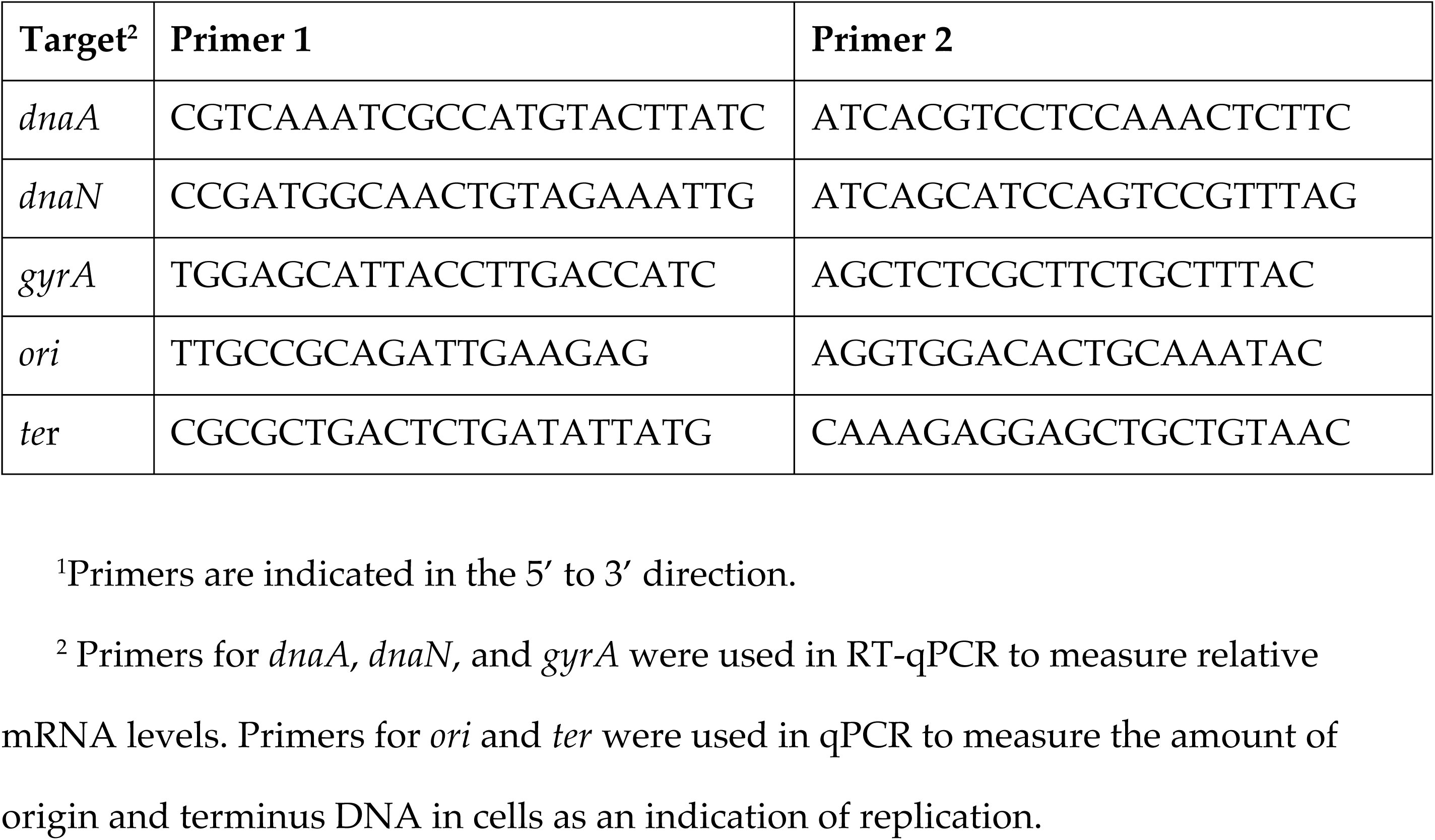
Primers for RT-qPCR and qPCR^1^.

**S2 Table.**
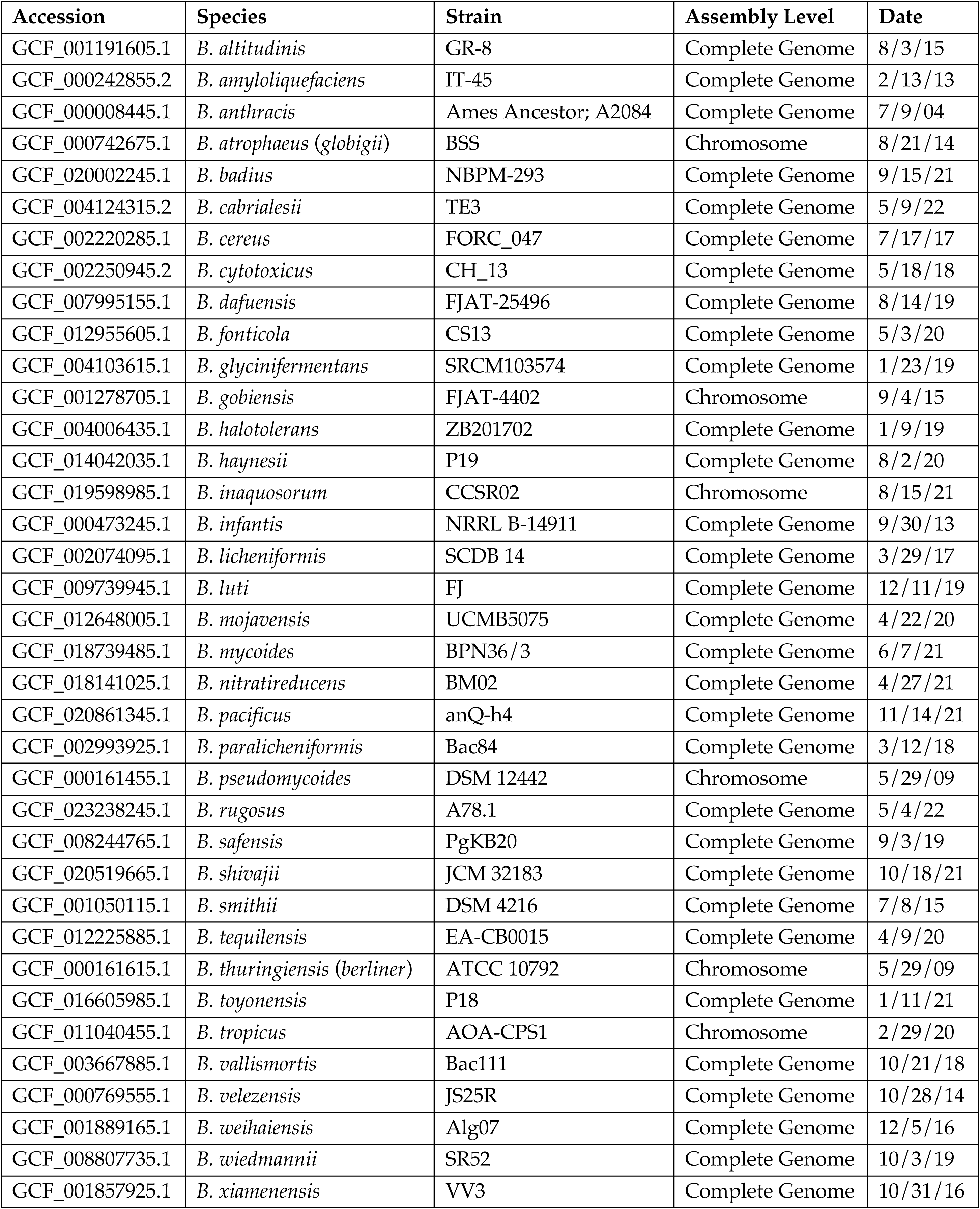
Reference genomes.

**S1 Data.** Underlying raw data. The excel spreadsheet contains the underlying data for experiments presented in the indicated figures and Table 1.

**S1 Raw Images.** Images of gels and blots used in Figure 4 and Figure 5. **S1A.** Images of the Northern blot (left) and stained gel (right) that were used for Figure 4A and 4B, respectively. **S1B.** Images of the Western blot (top) and stained gel (bottom) that were used for Figure 5B.

